# Integrative description of a new species of *Testechiniscus* (Tardigrada, Echiniscoidea) from Novaya Zemlya, Russia, with discussion of the genus diagnosis and distribution

**DOI:** 10.64898/2025.11.29.691272

**Authors:** Alexandra Yu. Tsvetkova, Denis V. Tumanov

## Abstract

In this paper we describe *Testechiniscus* **sp. nov.**, a new tardigrade species from Novaya Zemlya, Russia, using data from morphological and morphometric analyses conducted with the use of light and scanning electron microscopy, as well as from genetic analysis based on four molecular markers (three nuclear: 18S rRNA, 28S rRNA, ITS-1, and one mitochondrial: COI). A comprehensive differential diagnosis is provided. Upon examination of both new and the type species we describe internal leg plates as a new character for the genus *Testechiniscus*, amending the genus diagnosis. This study introduces new data on the geographic distribution of *Testechiniscus*.

## INTRODUCTION

Tardigrades are a group of microscopic segmented animals that are known for their ability to withstand extreme conditions in cryptobiotic state. They inhabit a wide range of aquatic biotopes from abyssal depths of the ocean to mountain tops (Nelson et al. 2018). The phylum currently comprises around 1500 species (Degma & Guidetti 2025).

In recent years the implementation of modern methods in tardigrade taxonomy revealed hidden diversity in most tardigrade taxa. The paradigm of widely distributed polymorphous species is being replaced with the concept of numerous local species, often poorly morphologically differentiated but clearly separated through molecular analysis (Bertolani et al. 2011; Gąsiorek et al. 2016, 2018a; Guidetti et al. 2019). Cosmopolitan nature of most species is now heavily scrutinized and most well-studied species show limited distribution (Gąsiorek et al. 2019a). Additionally, there are several species endemic to high alpine regions, as well as cases of split arctic-alpine habitats (Gąsiorek 2023a).

The genus *Echiniscus* C.A.S. Schultze, 1840 is one of the most speciose tardigrade genera and one of the earliest to be established. It is also long hypothesized to be polyphyletic; indeed, numerous molecular analyses and reconsiderations of morphological characters have resulted in isolation of numerous new genera preciously thought to be species groups within *Echiniscus* (Gąsiorek et al. 2018c; 2019b).

One such genus is *Testechiniscus* Kristensen, 1987, previously known as *spitsbergensis*-group (named after *Echiniscus spitsbergensis* Scourfield, 1897) and isolated on the basis of black crystalline eyes (*Echiniscus* species possess red eyes) and heavily sclerotized ventral surface of the body (Kristensen 1987). The original composition of the genus included *Testechiniscus spitsbergensis* as the type species, as well as three other species: *T. spinuloides* (Murray, 1907) (formerly *Echiniscus spinuloides*), *T. meridionalis* (Murray, 1906) (formerly *Echiniscus meridionalis*) and *T. clavisetosus* (Mihelčič, 1958) (formerly *Echiniscus clavisetosus*). Kristensen (1987) also noted the polar and alpine distribution of the newly established genus. Later, McInnes (1994) additionally assigned *Echiniscus macronyx* Richters, 1907 to the genus on the basis of ventral sculpturing and cephalic appendage configuration.

Gąsiorek et al. (2018b) reexamined the genus, redescribed the type species *T. spitsbergensis* from its terra typica (Spitsbergen), along with describing a subspecies *T. spitsbergensis tropicalis* Gąsiorek, Stec, Zawierucha, Kristensen & Michalczyk, 2018 from a high alpine biotope in Ethiopia. *Testechiniscus spinuloides* and *T. clavisetosus* have been dismissed as synonyms to *Testechiniscus spitsbergensis*. The authors also examined morphological characteristics of *T. meridionalis* and *T. macronyx* and concluded that both of these species deviate significantly from the morphological *Testechiniscus* diagnosis. Gąsiorek (2023b) reexamined morphological and molecular characters of *T. meridionalis* and restored its place within *Echiniscus*. Since *Echiniscus meridionalis* also possesses rows of ventral cuticular plates, this character can no longer be used as a criterion for distinguishing between *Testechiniscus* and *Echiniscus*. The reassessment of *T. macronyx* is still in progress and it is considered *Testechiniscus* sensu lato; its morphology does not resemble any echiniscid genera, which might lead to the establishment of a new genus within the family (Gąsiorek 2023b).

*Testechiniscus laterculus* (Schuster, Grigarick and Toftner, 1980) was initially recorded from North America by Grigarick et al. (1975) as *Echiniscus oihonnae* Schuster & Grigarick, 1965. Schuster et al. (1980) reexamined the “American” *E. oihonnae* and concluded that its morphology did not match the description of the species from European records. The authors chose to establish a new species of *Echiniscus* with the type locality in North America and renamed it to *Echiniscus laterculus*.

Interestingly, Kristensen (1987) did not include *E. laterculus* in the newly established genus *Testechiniscus*. Moreover, the article in question does not contain a single mention of the species (neither as *E. oihonnae* nor as *E. laterculus*). However, in the same year Dastych (1987) reexamined the species, along with several others within Echiniscidae, and confirmed the presence of ventral plates, which placed *E. laterculus* into the *spitsbergensis*-group. In the light of Kristensen’s recent work Dastych (1987) concluded that the species belonged to newly established genus *Testechiniscus*. However, an official reassignment was never given. In every relevant paper following Dastych’s reexamination (Kathman & Dastych 1990; McInnes 1994; Gąsiorek et al. 2018b) the species is referred to as *Testechiniscus laterculus*. Finally, Massa et al. (2024) obtained sequences of 18S and COI markers from a specimen of *T. laterculus* from Canada, making the description integrative; both sequences were used for molecular analysis in this study.

Currently only two species can be considered *Testechiniscus* sensu stricto: *T. spitsbergensis* (with 2 subspecies) and *T. laterculus* (Gąsiorek 2023b).

In this work, we describe a new species of the genus *Testechiniscus* from Novaya Zemlya combining modern molecular techniques with classical morphometric and morphological methods in an integrative approach.

## MATERIAL AND METHODS

### Sample processing

The moss and soil sample containing the new species was collected from Novaya Zemlya archipelago by Arseniy Kudikov on the 26th of September 2022. The material was dried and stored within a paper envelope at room temperature. Tardigrades were extracted from samples by washing them through two sieves (Tumanov 2018). The content of the fine sieve was examined under a Leica M205C stereo microscope. All animals extracted from the sample were divided into three groups for further analysis: light microscopy (LM) including phase contrast (PhC) and differential-interference contrast (DIC), scanning electron microscopy and DNA sequencing.

### Additional material

We examined the material of *T. spitsbergensis* from the SPbU collection (Department of Invertebrate Zoology, Faculty of Biology, St. Petersburg State University). The specimens were collected from Krasnoyarskiy kray, Sakhalin and Ireland (see Table 1). The specimen from Ireland was examined through a light microscope, whereas for *T. spitsbergensis* from Krasnoyarskiy kray and Sakhalin high-quality LM and SEM images were obtained. The original descriptions, as well as the revision of *T. spitsbergensis* and *T. laterculus* were used for comparison as well.

**Table 1.** Additional material of *Testechiniscus spitsbergensis* examined in the study.

| Locality | Substrate | Collector | Collection date | Light microscopy | SEM | Molecular data |
| --- | --- | --- | --- | --- | --- | --- |
| Krasnoyarskiy kray, Russia | Moss | Boris Kochnev | 02.08.2020 | + | + | + |
| Sakhalin, Russia | Moss/rock | Denis Tumanov | 19.08.2024 | + | + | + |
| County Clare, Ireland | Moss | Vadim Panov | 24.09.2004 | + | – | – |

### Microscopy and imaging

Specimens for light microscopy (LM) were fixed with acetic acid or relaxed by incubating live individuals at 60°C for 30 min (Morek et al. 2016). Additionally, several specimens were placed into a mix of glycerol, ethanol and water in proportion of 1:1:1, and kept at room temperature until the complete evaporation of ethanol and water. Afterwards the animals were mounted on slides in Hoyer’s medium.

Permanent slides were examined under Leica DM2500 microscope equipped with phase contrast (PhC) and differential interference contrast (DIC), associated with a Nikon DS-Fi3 digital camera and NIS software.

As preparation for scanning electron microscopy tardigrades were subjected to a 60°C bath (Morek et al. 2016), then dehydrated in a series of water-ethanol mixtures (10%, 20%, 30%, 50%, 70%, 96% ethanol), an ethanol-acetone 1:1 mixture, and 100% acetone. The specimens then underwent CO_2_ critical point drying and after that they were placed on stubs and sputter coated with a thin layer of gold or platinum (50 nm). Ultrastructural analysis of the specimens’ body surface was conducted under high vacuum in a Tescan mIRA3 Lmu Scanning Electron microscope at the Centre for Molecular and Cell Technologies, St Petersburg State University and Hitachi TM-1000 (Hitachi, Japan) at “Taxon” Research Resource Center of the Zoological Institute of the Russian Academy of Sciences.

All figures were assembled in Adobe InDesign CS4. For structures that could not be satisfactorily focused in a single LM photograph, a stack of 2–6 images was taken and assembled manually into a single deep-focus image, using Helicon Focus 6. Panorama maker 6 was used to combine a series of images in cases where the whole structure didn’t fit into the camera field of view under high magnification.

### Morphometrics and morphological nomenclature

Specimens were measured under LM using phase contrast, associated with a Nikon DS-Fi3 digital camera and NIS software. Sample size was adjusted following recommendations by Stec et al. (2016). All measurements are given in micrometers (μm). Structures were measured only if their orientation was suitable. Body length was measured from the anterior extremity to the end of the body, excluding the hind legs. The *sc* ratio is the ratio of the length of a given structure to the length of the scapular plate (Fontoura & Morais 2011). Ventral armature denotation system follows Kaczmarek et al. (2012). Body and trunk appendage configuration is given according to Ramazzotti & Maucci (1983) and amendments by Gąsiorek et al. (2017). Leg morphology is described following Kristensen (1987), external leg plate configuration is given according to DeMilio et al. (2022). Morphometric data were handled using the Echiniscoidea ver. 1.2 template from the Tardigrada Register, www.tardigrada.net/register (Michalczyk & Kaczmarek 2013).

### Genotyping

DNA was extracted from individual specimens using QuickExtract™ DNA Extraction Solution (Lucigen Corporation, USA, see complete protocol description in Tumanov 2020). Due to an extraordinarily thick cuticle of *Testechiniscus* species first attempts at retrieving DNA from intact specimens were unsuccessful. Further attempts included piercing the specimens with a sterile thin needle before placing them into the solution, which yielded better results. Preserved exoskeletons were recovered, mounted on a microscope slide in Hoyer’s medium and retained as the hologenophore (Pleijel et al. 2008).

Four genes were sequenced: a small ribosomal subunit (18S rRNA) gene, a large ribosomal subunit (28S rRNA) gene, internal transcribed spacer (ITS-1) and the cytochrome oxidase subunit I (COI) gene. PCR reactions included 5 μl template DNA, 1 μl of each primer, 1 μl DNTP, 5 μl Taq Buffer (10X) (−Mg), 4 μl 25 mM MgCl_2_ and 0.2 μl Taq DNA Polymerase (Thermo Scientific™) in a final volume of 50 μl. The primers and PCR programs used are listed in Supplementary Material 01. The PCR products were visualized in 1.5% agarose gel stained with ethidium bromide. All amplicons were sequenced directly using the ABI PRISM Big Dye Terminator Cycle Sequencing Kit (Applied Biosystems) with the help of an ABI Prism 310 Genetic Analyzer in the Core Facilities Center ‘Centre for Molecular and Cell Technologies’ of St. Petersburg State University. Sequences were edited and assembled using ChromasPro software (Technelysium). The COI sequences were translated to amino acids using the invertebrate mitochondrial code, implemented in MEGA11 (Tamura et al. 2021), in order to check for the presence of stop codons and therefore of pseudogenes. GenBank accession numbers of all sequences used in the analysis are given in Supplementary Material 02.

## RESULTS

### Reexamination of *T. spitsbergensis* morphology

We examined a population of *T. spitsbergensis* from Krasnoyarskiy kray (Fig. 1) and another one from Sakhalin, comparing them with the species description given in Gąsiorek et al. (2018b). Several morphological characters were reexamined that call for adjusting that description, as well as the genus diagnosis.

**Fig. 1.**
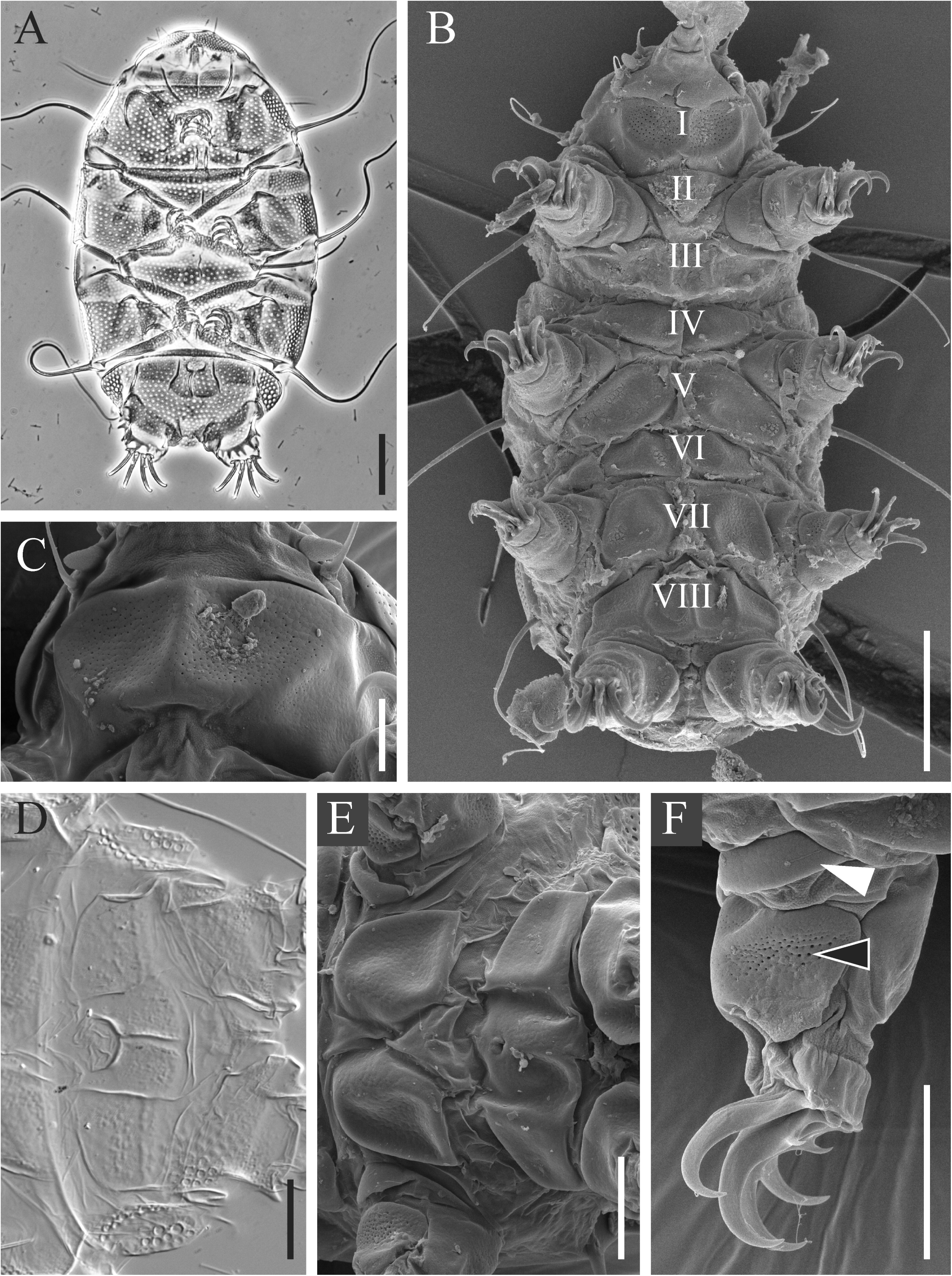
*Testechiniscus spitsbergensis*, details of morphology. **A.** Habitus, dorso-ventral view, PhC. **B.** Ventral view, SEM. **C.** Subcephalic plate, SEM. **D.** Genital plate II (row VIII), DIC. **E.** Genital plates I and II (rows VII-VIII), SEM. **F.** Internal leg plates on leg I, SEM. Internal coxal plate is marked by a white arrowhead, internal femoral plate indicated by a black arrowhead. Scale bars: **A, B** = 50 μm, **C** = 10 μm, **D, E, F** = 20 μm.

#### Ventral plates (Fig. 1B–E)

We interpret a plate as continuous when its posterior edge is unbroken and there are no areas of soft cuticle between the sections. Seams or ridges that are structurally part of the plate are not interpreted as a divide between two paired plates but instead a visual divide of an unpaired one. Based on this understanding, we reexamined the ventral plates in *T. spitsbergensis* from both populations.

Subcephalic plate (row I) is very large, covering the entire underside of the head. Well-developed and visible under both LM and SEM, it consists of two sections covered in large pores with a ridge in the middle. Under SEM, it is clear that the posterior edge of the subcephalic plate is continuous (Fig. 1C). The ridge is part of the plate itself, and there are no areas of flexible cuticle dividing the sections. We therefore propose describing the subcephalic plate in *T. spitsbergensis* as unpaired.

Genital plate II (row VIII) in *T. spitsbergensis* from both localities bears four sections of approximately equal size that are visible both in LM and SEM (Fig. 1D, E). Under LM, all four sections have distinct sculpturing, different from the surrounding soft ventral cuticle. Under SEM, the lateral sections of the plate are raised, while the medial sections are sunken into the body, appearing level with the rest of the cuticle. Sections are visually divided by three ridges, similar to that on the subcephalic plate. However, the cuticular sculpture on the ridges is not drastically different from that on the sections themselves. Moreover, there is a distinct posterior edge that is unbroken for the entire row VIII (Fig. 1E). Additionally, areas of soft cuticle, present between truly divided paired plates, are absent here. All of the above serves as a clear indication of row VIII being comprised of a single continuous plate.

Based on our findings, the amended ventral plate formula we propose for *T. spitsbergensis* is VIII: 1-1-2-2-2-6-2-1.

#### Internal leg plates (Fig. 1F)

In addition to the ventral plates, very well-defined internal plates on legs I–IV are present in both populations of *T. spitsbergensis*. Legs I–III bear 2 small plates each – coxal and femoral. Internal plates on legs IV are very large, completely covering the internal surface of the leg and wrapping around the sides (Fig. 1B). Such plates on leg IV is a character never before described for the genus *Testechiniscus*. Internal coxal and femoral plates on legs I–III are shifted to the anterior side of the leg. The femoral internal-external plate pair constitutes a continuous structure wrapping around the leg, whereas the coxal plates are separate with an area of flexible cuticle between them.

### Description of the new species

#### Taxonomic account of the new species

Phylum: Tardigrada Doyère, 1840

Class: Heterotardigrada Marcus, 1927

Order: Echiniscoidea Richters, 1926

Family: Echiniscidae Thulin, 1928

Genus: *Testechiniscus* Kristensen, 1987

Species: *Testechiniscus* **sp. nov.**

#### Candidates for the type series

Slides 314(3)–314(14), 314(21), 314(25)–314(30), 314(32)–314(40), 314(42)–314(63), 314(65) from Novaya Zemlya archipelago, Russia; 20 specimens on 3 SEM stubs are deposited in SPbU (Stubs 63, 69, 83)).

#### Type locality

Russia; Novaya Zemlya archipelago, Severny Island, Blagopoluchiya Bay, N 75°44’ 7.27224’’ E 63°35’25.37628’’, moss and soil, 26 Sept. 2022, Arseniy Kudikov. Found together with *Ursulinius* cf. *elegans*.

#### Morphology

Adults (from the third instar onward, measurements and statistics in Table 2). Body bright orange (Fig. 2C), with large black eyes that stay visible after mounting (Fig. 3F). An *Echiniscus*-type cephalic papilla and two cirri on well-developed cylindrical cirrophores. Small primary clava very close to the cirrophore A (Fig. 3A, B). The body appendage configuration in adults is *A-(B)-B^l^-C-C^l^-C^d^-D-D^l^-D^d^*, of which dorsolateral appendages are always short spines (Fig. 2A, B). Cirri B often absent – in 8 out of 20 measured specimens (40%). Additionally, 1–3 small spines may be present along notches of the caudal (terminal) plate (Fig. 3C, D, E).

**Fig. 2.**
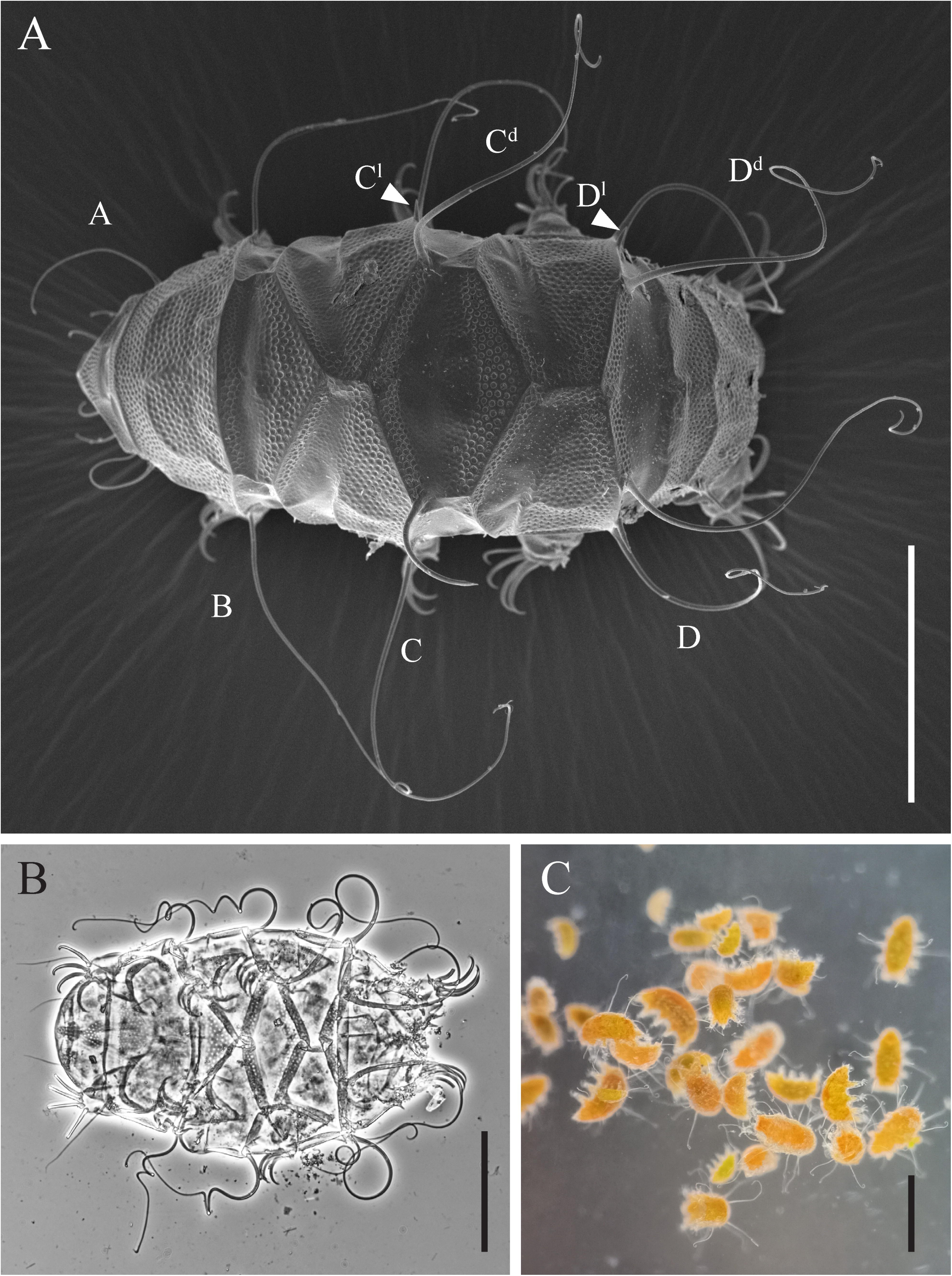
*Testechiniscus* **sp. nov.**, habitus. **A.** Paratype, SPbU_69, dorsal view, SEM. Dorsolateral appendages are marked by white arrowheads. **B.** Paratype, 314(26), dorso-ventral view, PhC. **C.** Coloration in living specimens under incident light. Scale bars: **A, B** = 100 μm, **C** = 300 μm.

**Fig. 3.**
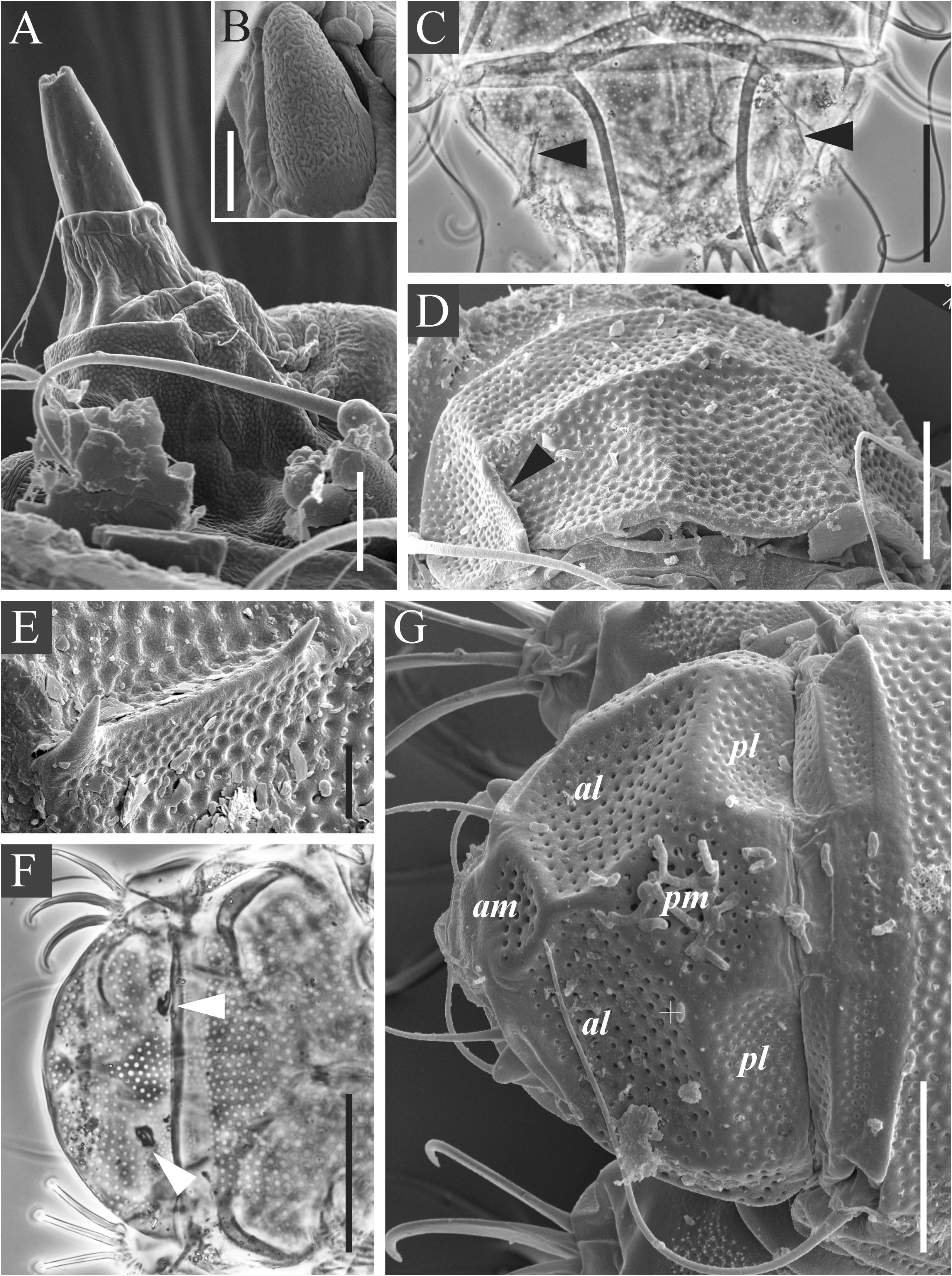
*Testechiniscus* **sp. nov.**, details of cuticular structures. **A, B.** Paratype, SPbU_63. **C, F.** Paratype, 314(26). **D, G.** Paratypes, SPbU_83. **E.** Paratype, SPbU_69. **A.** Side view of the buccal cone, granular ventral sculpture on the head, SEM. **B.** Clava, SEM. **C.** Caudal plate, PhC. **D.** Caudal plate, SEM. Notches of the caudal plate are indicated by black arrowheads. **E.** Spines along the notch of the caudal plate, SEM. **F.** Anterior end of the body, dorso-ventral view, PhC. Eyes are marked by white arrowheads. **G.** Cephalic and cervical (neck) plate, SEM. Subdivisions of the cephalic plate are indicated in italics. Scale bars: **A, E** = 5 μm, **B** = 2 μm, **C, F** = 50 μm, **D, G** = 20 μm.

**Table 2.**
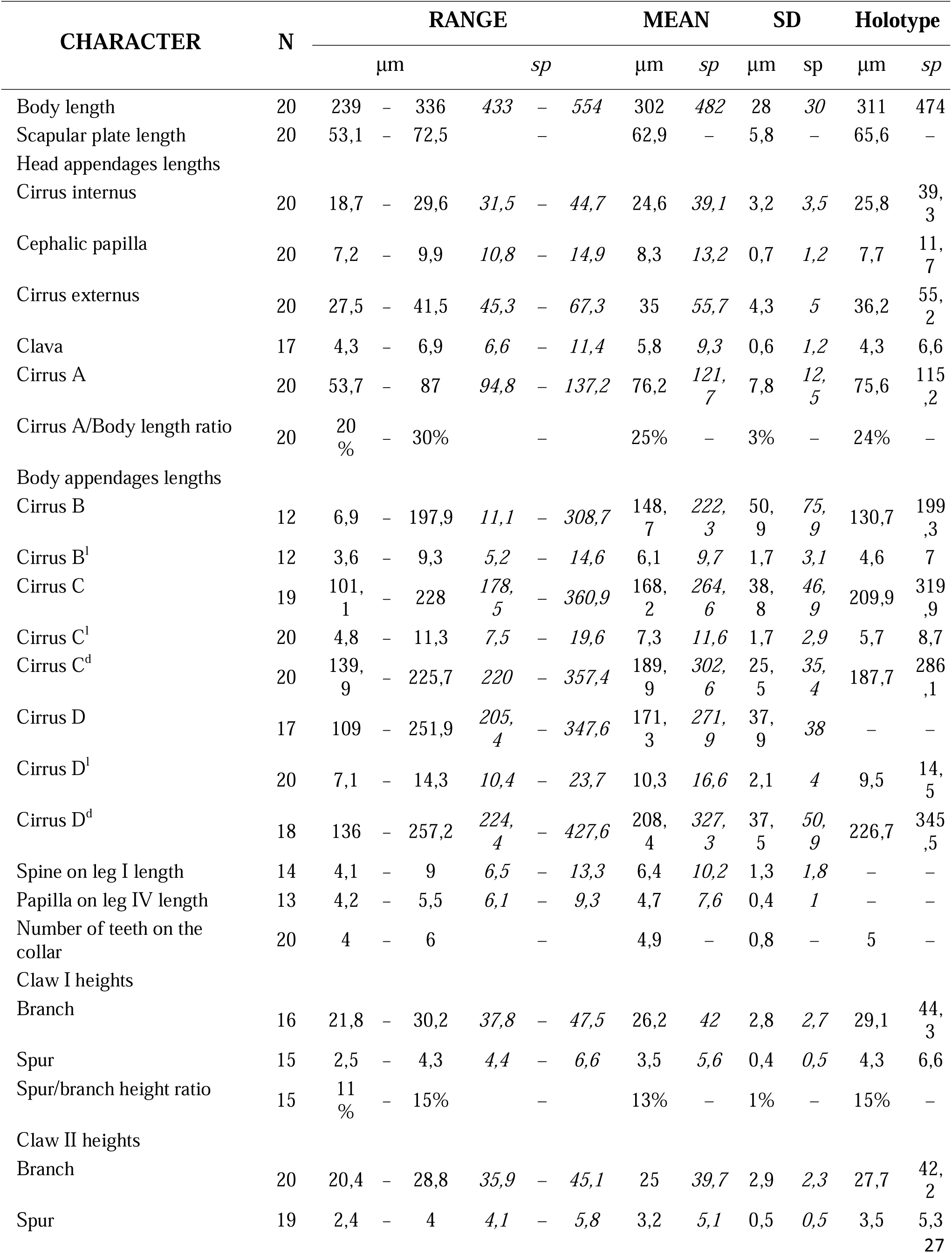
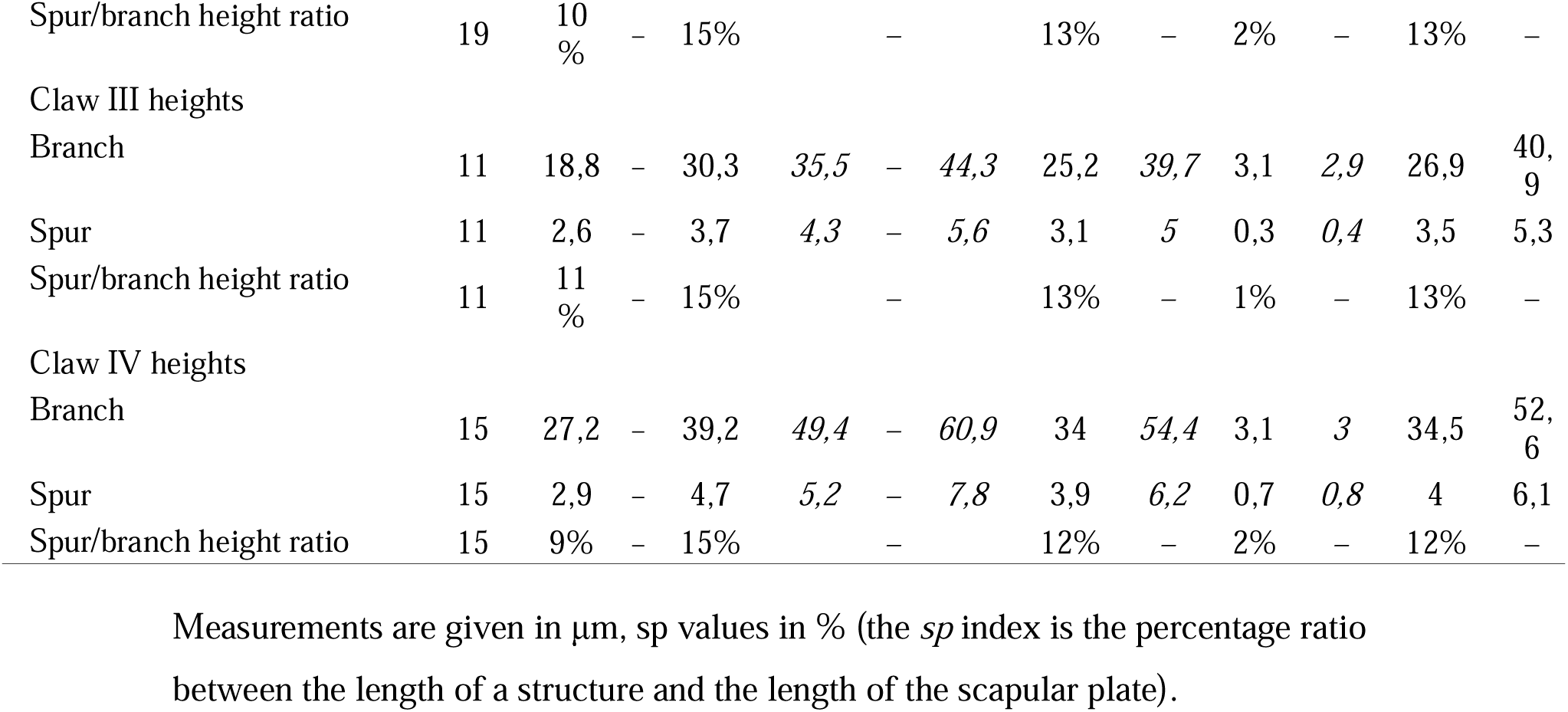
Measurements [in μm] of selected morphological structures of adult females (the 3rd and older instars) of *Testechiniscus* sp. nov.

Dorsal plates with the *spitsbergensis* type sculpturing (originally the *blumi-canadensis* type): large, densely arranged, round true pores on the scapular, median I, caudal and posterior portions of median II and segmental plates (Fig. 4A–E). The true pores are surrounded by intracuticular cavities which appear as polygonal shapes in LM but are not visible under SEM (Fig. 4A). Anterior portions of median II and paired segmental plates, and the entire median III covered with evident reticulum (Fig. 4D–G).

**Fig. 4.**
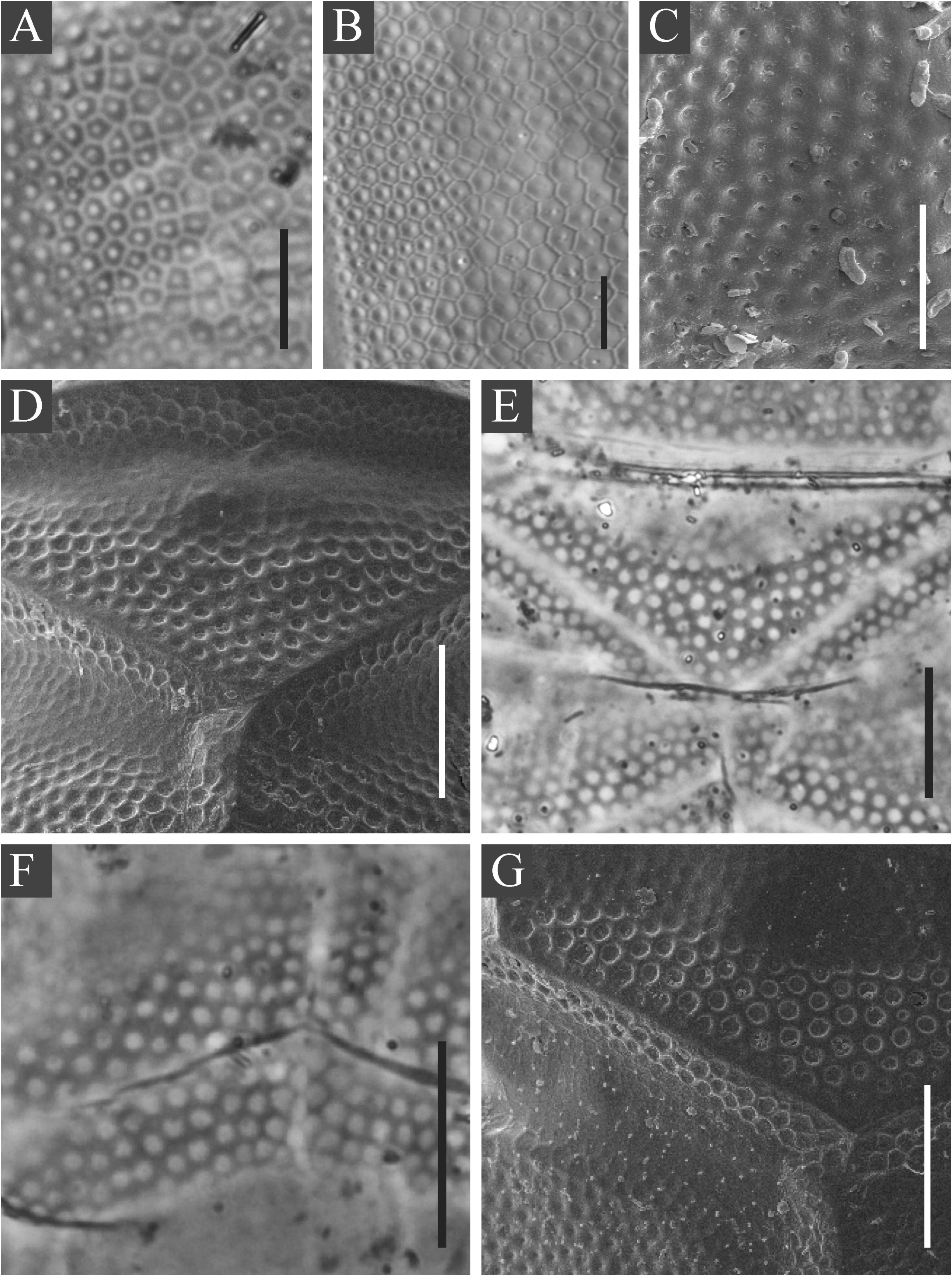
*Testechiniscus* **sp. nov.**, cuticular sculpture. **A, E, F.** Paratype, 314(7)-2. **B.** Paratype, 314(26). **C.** Paratype, SPbU_83. **D, G.** Paratype, SPbU_69. **A.** Scapular plate sculpture, PhC. **B.** Scapular plate sculpture, DIC. **C.** Scapular plate sculpture, SEM. **D.** Median plate 1, SEM. **E.** Median plate 1, PhC. **F.** Posterior edges of paired segmental plates II and anterior edge of the caudal plate, PhC. **G.** Posterior edge of median plate 2 and anterior edges of paired segmental plates II, SEM. Scale bars: **A**, **B**, С = 10 μm, **D**, **E**, **F**, **G** = 20 μm.

Cephalic plate well-developed, consisting of six prominent sections separated by ridges (Fig. 3G). For a proposed new nomenclature system for these subdivisions see Discussion. The sections are covered with pores whereas the ridges are smooth or with strongly reduced pores (Fig. 3F, G). Cervical (neck) plate rectangular, with a transversal ridge separating it into two sections. Both sections are covered with pores (Fig. 3G). Scapular plate very large. Paired segmental plates divided into a narrower anterior and a larger posterior part, with an indistinct border between them, i.e. pores covering the posterior part become shallow and gradually convert into reticulum on the anterior part (Figs 3A; 4E, G). The anterior margins of paired segmental plates are less sclerotized than the posterior. The caudal (terminal) plate slightly smaller than the scapular one, with paired notches (incisions) and a transverse ridge dividing it into two sections: a larger latero-frontal section and a smaller medio-caudal section (Fig. 3C, D). Both sections bear pores. Median plate I and median plate II divided into two unequal parts (Fig. 2A). Median plate III small, and often partly hidden under the posterior edges of segmental plate II and under the central portion of the caudal (terminal) plate (Fig. 2B). Lateral intersegmental plates absent.

Venter with 8 rows of well-defined plates of various sizes (Figs 5, 6). Subcephalic plate (row I) large, completely devoid of pores but bearing three ridges and weakly developed epicuticular elements (visible only under SEM as indistinct bumps) (Fig. 6A). Plates in rows II and V smooth (Fig. 6A, C). Row III consists of two median smooth plates and two lateral plates with distinct bumps visible under SEM (Fig. 6B). Rows IV and VI are constituted by a single unpaired plate with rugose sculpturing (Fig. 6B, C). Genital plates I (row VII) paired, also bear weak epicuticular elements visible under SEM. Genital plate II (row VIII) unpaired, with dimpled surface under SEM and pillars in the inner epicuticular layer visible in LM (Fig. 6D). Only one unpaired plate, however, is truly undivided (row II), whereas unpaired plates in rows I, IV, VI, and VIII are visually divided into two symmetrical halves by either a longitudinal ridge (rows I and VIII) or a seam-like structure that disrupts the surrounding epicuticular patterns (rows IV and VI). Ventral plate formula is VIII: 1-1-4-1-2-1-2-1. Posterior edge of unpaired ventral plates often extends over the body cuticle, forming an overhang visible under LM and SEM (Fig. 6E).

**Fig. 5.**
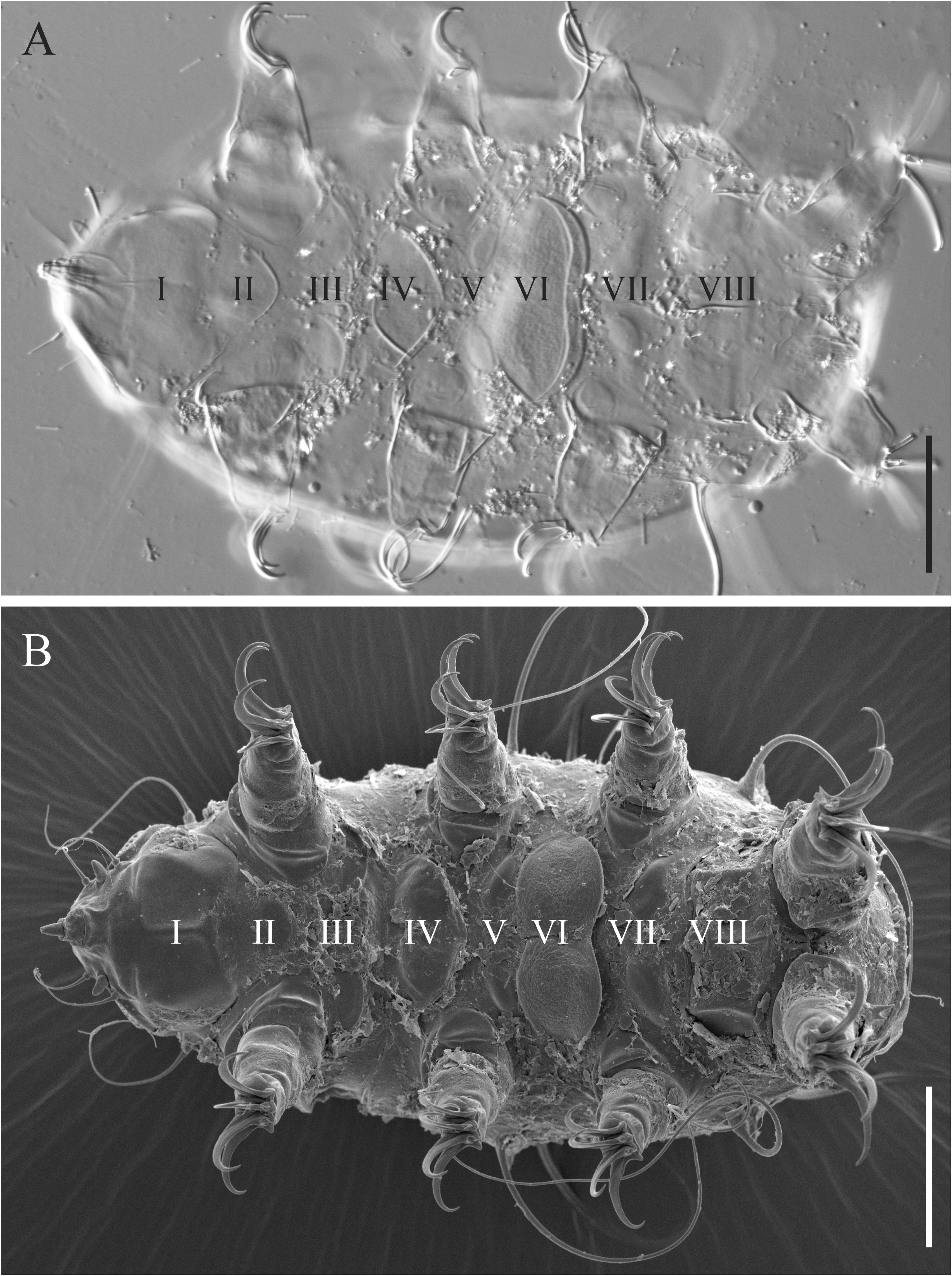
*Testechiniscus* **sp. nov.**, ventral plates. **A.** Paratype, 314(7)-2. **B.** Paratype, SPbU_69. **A.** Ventral view, DIC. **B.** Ventral view, SEM. Rows of ventral plates indicated by roman numerals. Scale bars = 50 μm.

**Fig. 6.**
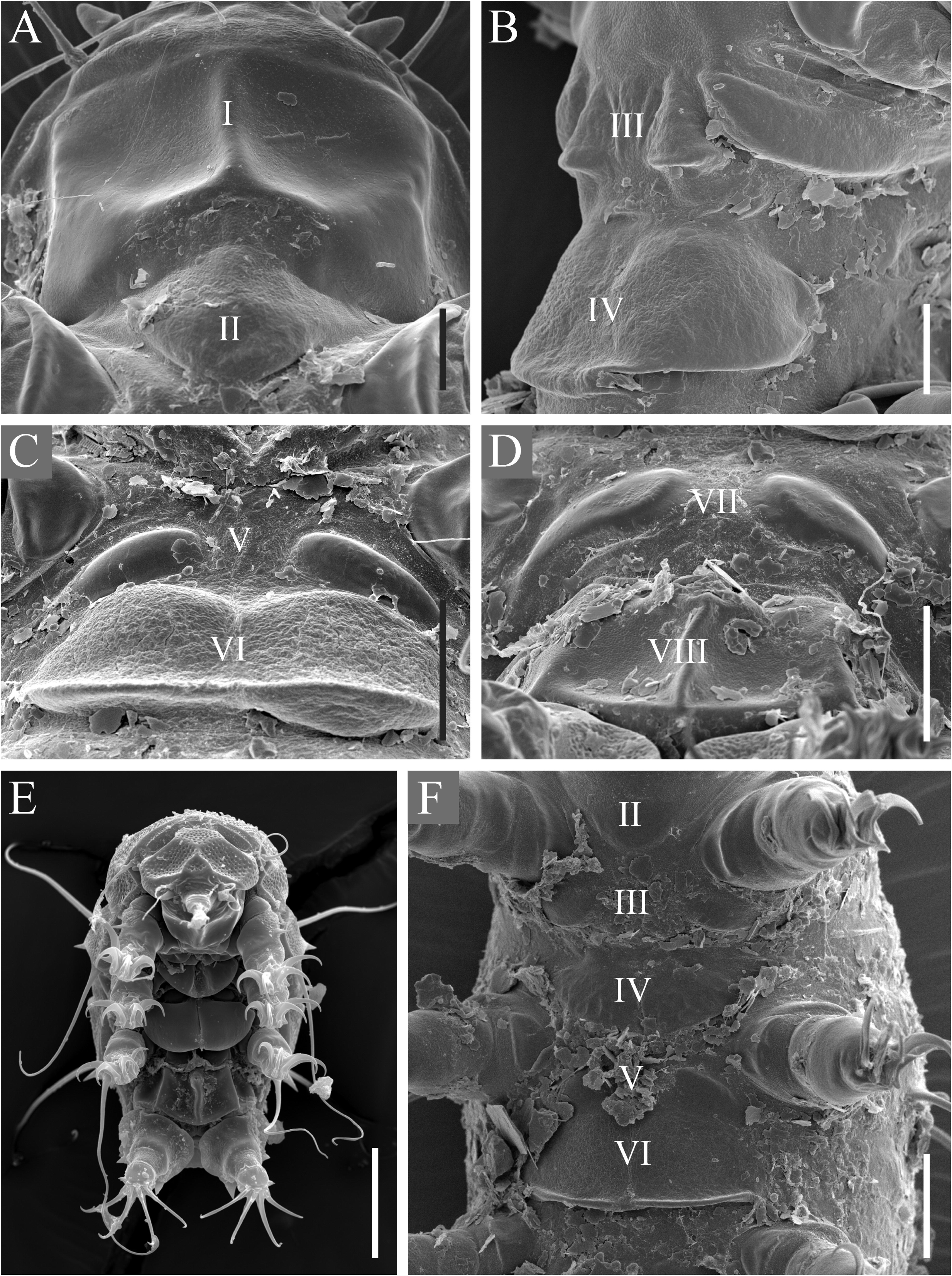
*Testechiniscus* **sp. nov.**, ventral plates (continued), SEM. **A, C, D.** Paratype, SPbU_69. **B.** Paratype, SPbU_63. **E, F.** Paratypes, SPbU_83. **A.** Rows I (subcephalic plate) and II. **B.** Rows III and IV. **C.** Rows V and VI. **D.** Rows VII (genital plate I) and VIII (genital plate II). **E.** Ventral plates protruding above the body surface. **F.** Underdeveloped ventral plates in a juvenile specimen. Scale bars: **A, B** = 10 μm, С**, D, F** = 20 μm, **E** = 50 μm.

Gonopore in females separated from anus, with a typical six-leaved rosette of cells surrounding the opening. No males were found in the examined material.

The new species possesses a set of pronounced coxal and femoral leg plates (Fig 7A–F). Femoral plates on legs I–III with distinct sculpture and bear pores, whereas coxal plates are often less developed and smooth. A large spine on legs I present (Fig. 7A, B); coxal plate on legs IV bears a very small papilla, often covered with substrate particles and barely (if at all) visible in LM (Fig. 7F). Dentate collar on legs IV with 4–6 irregular teeth (Fig. 7D, E). External claws on legs I–III smooth, secondary spurs on legs IV absent (Fig. 7G–J). Claw bases well-developed, often with sharpened processes (Fig. 7K). Internal claws with widened and concave portions between claw base and spur (Fig. 7L). Spurs on internal claws large, positioned at ca. 20–30% of the claw height. Spurs on legs I-III directed downwards (Fig 7G), whereas on leg IV they are positioned at an almost 90% angle relative to the branch (Fig 7I).

**Fig. 7.**
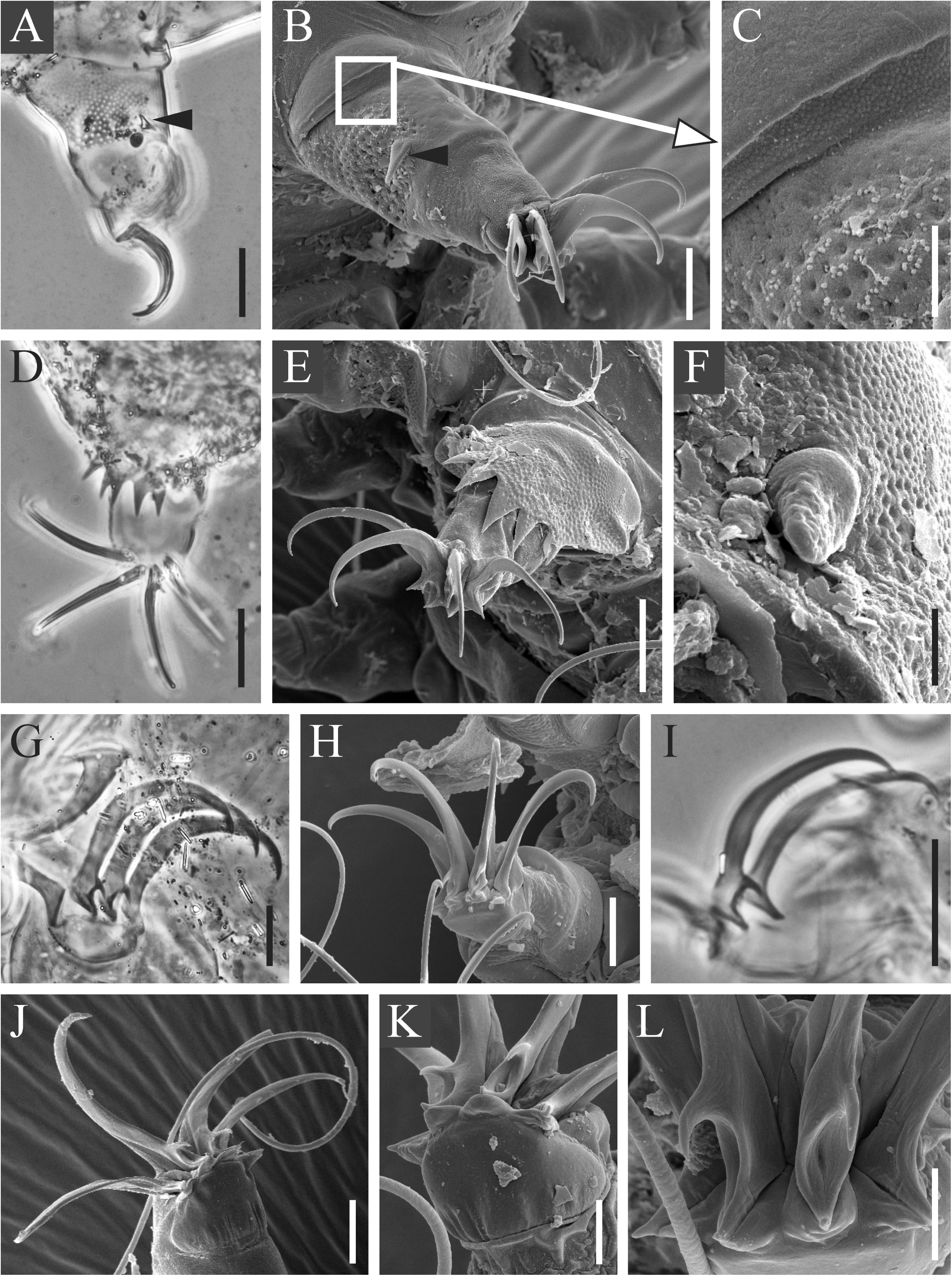
*Testechiniscus* **sp. nov.**, legs. **A, I.** Paratype, 314(8)-2. **B, C, E, F, J.** Paratype, SPbU_69. **D.** Paratype, 314(7)-2. **G.** Paratype, 314(26). **H**, **K, L.** Paratype, SPbU_83. **A.** Leg I, external view, PhC. **B.** Leg I, SEM. Black arrowheads indicate the spine on leg I. **C.** Fragment of leg I with edges of the coxal and femoral plates and an area of soft granulated cuticle between them, SEM. **D.** Leg IV, PhC. **E.** Leg IV, SEM. **F.** Papilla on leg IV, SEM. **G.** Claws on leg II, PhC. **H.** Claws on leg II, SEM. **I.** Claws on leg IV, PhC. **J.** Claws on leg IV, SEM. **K.** Claw bases with sharpened processes, SEM. **L.** Widened and concave portions between claw base and spur on internal claws, SEM. Scale bars: **A, D, E, I** = 20 μm, **B, G, H, J** = 10 μm, С = 4 μm, **F** = 2 μm, **K, L** = 5 μm.

The terminal part (tarsus) of the leg is telescopic, often retracting inward during SEM specimen preparation. In living specimens, the claws are mobile, packed together during the forward movement of the leg and spreading into a fan shape when the leg is moved backwards. The described movement is illustrated in the Supplementary Material 03.

Just like *T. spitsbergensis*, the new species possesses very well-defined internal plates on legs I–IV (Fig 8A–E). Plate configuration identical to that of the type species: coxal and femoral plates on legs I–III (Fig. 8A–C) and very large and wide internal plates on legs IV (Fig. 8D). Plates on legs I–III smooth except for internal femoral plates on leg III which demonstrate distinct dimpled surface under SEM. Identically to *T. spitsbergensis*, in *T.* **sp. nov.** the femoral plates connect anteriorly, whereas the coxal plates are separate (Fig. 8E).

**Fig. 8.**
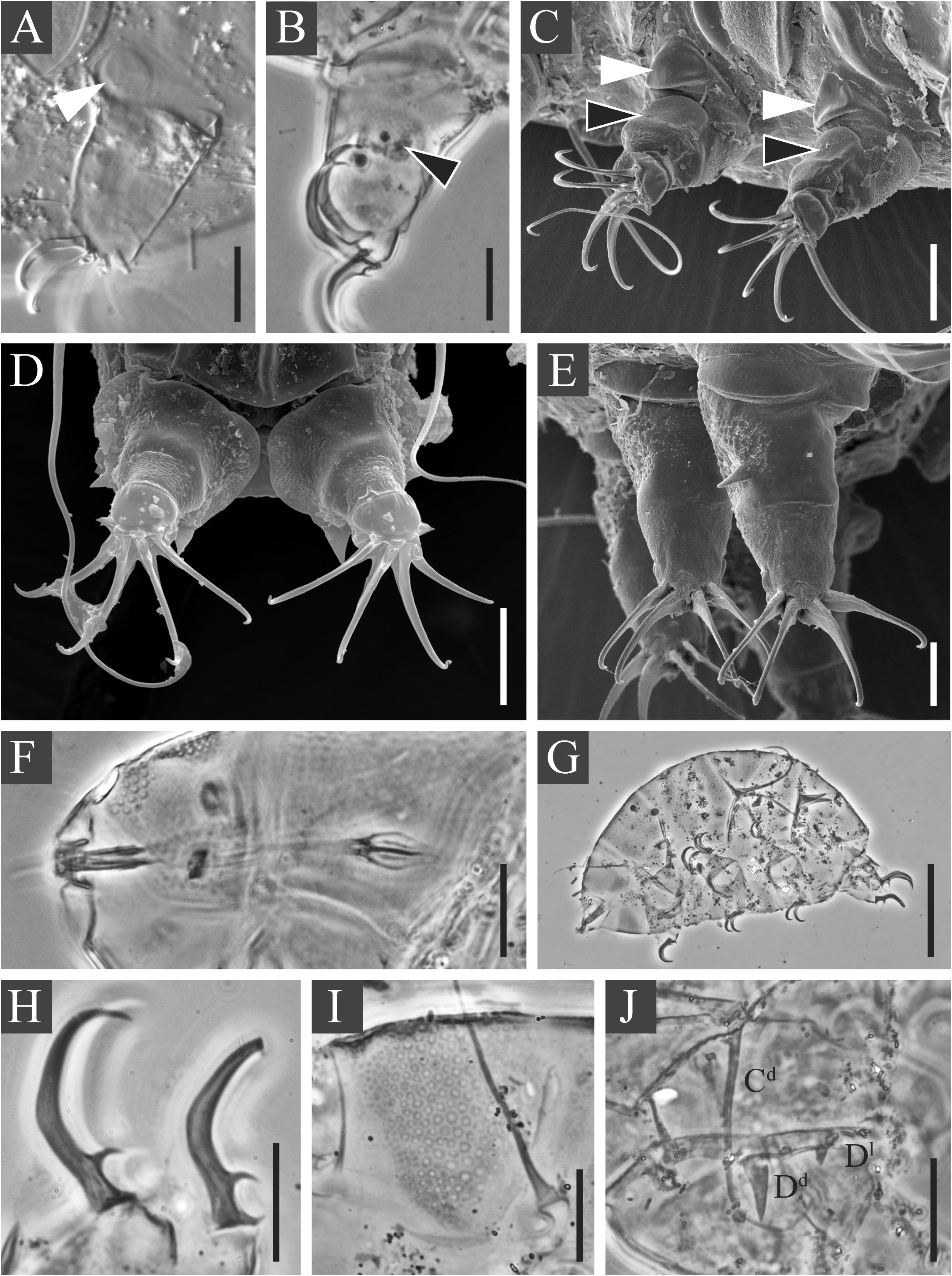
*Testechiniscus* **sp. nov. A.** Paratype, 314(7)-2. **B.** Paratype, 314(8)-2. **C, E.** Paratypes, SPbU_69. **D.** Paratype, SPbU_83. **F.** Paratype, 314(7)-1. **G, H, I.** Paratype (larva), 314(55). **J.** Paratype (larva), 314(54)-2. **A.** Internal view of leg III, DIC. **B.** Internal view of leg I, PhC. **C.** Internal view of legs II and III, SEM. Internal coxal plates are marked by white arrowheads, internal femoral plates indicated by black arrowheads. **D.** Large internal plates on legs IV, SEM. **E.** Anterior view of legs I and II, SEM. **F.** Bucco-pharyngeal apparatus, PhC. **G.** Larva, habitus (lateral view), PhC. **H.** Larva, claws on legs IV, PhC. **I.** Larva, scapular plate, PhC. **J.** Cirri in larva, PhC. Scale bars: **A, B, C, D, F, J** = 20 μm, **E, H, I** = 10 μm, **G** = 50 μm.

Buccal apparatus short, with rigid tube and roundish bulbus (Fig. 8F). Stylet supports large, cuticular, not encrusted with CaCO_3_ (see Kristensen 1987).

Juveniles (i.e. the second instar). Body bright orange. Morphology generally similar to that of adults; cirrus D is a short spine; no gonopore. Weakly developed ventral plates (Fig. 6F).

Larvae (i.e. the first instar). Body orange. Cirri D^d^ always long spines (appendage configuration: *A-C^d^-D^l^-D^d^*). Pores in the dorsal plates well-developed. Pedal (leg) plates are not visible under PCM. No anus and gonopore, only internal claws are present (Fig. 8G–J).

Dorsum and venter mucous, typically with attached substrate particles (Figs 3A, G; 5B). The level of cuticle mucousity corresponds to that described for the genus (Gąsiorek et al. 2018).

#### Molecular markers

We obtained good quality sequences for the 18S rRNA, 28S rRNA, COI and ITS-1 fragments, but we failed to amplify the ITS-2 marker. All sequenced fragments represented by single haplotypes (for full list of sequences see Supplementary Material 02).

### Differential diagnosis

#### Phenotypic

*Testechiniscus* **sp. nov.** can be distinguished from *T. spitsbergensis* by the ventral plate configuration (VIII: 1-1-4-1-2-1-2-1 in the new species vs VIII: 1-1-2-2-2-6-2-1 in *T. spitsbergensis*) and by the structure of the subcephalic plate (epicuticular bumps on the surface, pores absent in the new species, pores present in *T. spitsbergensis*).

*Testechiniscus* **sp. nov.** can be distinguished from *T. laterculus* by: the appendage configuration (*A-(B)-B^l^-C-C^l^-C^d^-D-D^l^-D^d^* in the new species vs *A-C-C^l^-D-D^l^-D^d^-E* in *T. laterculus*), the ventral plate configuration (VIII: 1-1-4-1-2-1-2-1 in the new species vs VIII: 2-1-2-1-2-1-2-2 in *T. laterculus*), by the presence of well-defined inner pedal plates on all legs, and by cuticular sculpturing (large pores in the new species vs small pores in *T. laterculus*).

#### Genotypic

The *p*-distances between *T. spitsbergensis* and *T.* **sp. nov.** were as follows: 18S rRNA: 0.22–0.34%, 28S rRNA: 0.46–0.62%, ITS-1: 3.14–3.47%, COI: 14.53–14.91%.

The *p*-distances between *T. laterculus* and *T.* **sp. nov.** were: 18S rRNA: 1.01%, COI: 15.30%.

## DISCUSSION

### Proposing a new nomenclature for cephalic plate sections

A definitive designation system for the *Testechiniscus* cephalic plate divisions has not yet been established. Previous descriptions of this plate either mention the faceted structure without specification (Kristensen 1987) or contain descriptions of five facets, which are, however, not numbered or otherwise individually indicated in illustrative material (Gąsiorek et al. 2018b). Upon examining SEM images of *T. spitsbergensis spitsbergensis* and *T. spitsbergensis tropicalis* provided by the authors the cephalic plates of both subspecies appear to be divided into six facets instead of five. Moreover, the new species also possesses a cephalic plate with six sections that are visible both under SEM and LM. The lack of definitive nomenclature, therefore, may create confusion in any future taxonomic investigations regarding the genus.

We propose a new system of names for the cephalic plate subdivisions as follows: ***am*** for anterior medial section (1), ***al*** for anterior lateral sections (2), ***pm*** for posterior medial section (1), and ***pl*** for posterior lateral sections (2) (Fig. 3G).

### Amended description of the type species *T. spitsbergensis*

As stated in Results, upon reexamination of the type species’ morphology we suggest changing the ventral plate formula for *T. spitsbergensis* to VIII: 1-1-2-2-2-6-2-1.

### Introducing internal leg plates as a new character for the genus *Testechiniscus*

As stated in the description, the new species possess well-developed cuticular plates not only on the outer surface of the legs but also on the inner surface of all legs (I–IV). The inner pedal plates are very pronounced in adults and juveniles but are not visible in larvae under PCM. Interestingly, this paper is not the first time such a character has been noted for the genus: in his work dedicated to the revision of Echiniscidae and establishing, among others, the new genus *Testechiniscus*, Kristensen (1987) states in the description of the genus morphology: “At least the type species *T. spitsbergensis* has attitudinal leg plates on the inner side of the three first legs.” However, these plates have not been given a detailed description by the author. Moreover, in the revision of *Testechiniscus* Gąsiorek et al. (2018b) did not mention such a character at all, both in the genus diagnosis and in the integrative redescription of the type species.

Upon reexamination of *T. spitsbergensis* from Krasnoyarskiy kray and Sakhalin we can conclude that the type species also possesses internal leg plates, visible both under PCM and SEM. The undoubtable presence of these structures in several species of *Testechiniscus* calls for the inclusion of the rediscovered character into the morphological diagnosis of the genus.

### Amended diagnosis of *Testechiniscus* (based on the diagnosis in Gąsiorek et al. 2018b)

Medium-sized echiniscids possessing black crystalline eyes. Rigid buccal tube with large cuticular stylet supports. Appendaged, i.e. having both cephalic and trunk cirri. Two pairs of segmental plates, unpaired scapular and caudal plates. Three median plates. Cuticular sculpture composed of large true round or polygonal pores that gradually become reticulum. Incisions (notches) on caudal (terminal) plate.

Eight rows of ventral plates. No pseudosegmental plates. All legs with a set of external plates (coxal and femoral) and internal plates (coxal and femoral). Femoral plates of each external-internal pair often form a continuous structure wrapping around the anterior side of the leg.

### Geographic distribution of the genus in Russia and worldwide

Molecular data confirms the presence of *T. spitsbergensis* in Krasnoyarskiy kray and on Sakhalin (see Supplementary Material 02). Other records are exclusively morphological – the species has been reported from Caucasus (Biserov 1991), Taymyr Peninsula and Novaya Zemlya archipelago (Biserov 1996, 1999), as well as from Franz-Josef Land (Murray 1907; Richters 1911). *Testechiniscus laterculus* has also been recorded from Taymyr (Biserov 1996) but molecular data is needed for verifying these reports. Initially, the description of *T.* **sp. nov.** from Novaya Zemlya supported arcto-alpine distribution for the genus on Russian territory, with all species located in Russia being restricted to the high arctic and mountainous regions. However, a population of *T. spitsbergensis* that was recently extracted from moss on rocks above the tidal zone on the southern coast of Sakhalin breaks this pattern, adding a temperate non-mountainous habitat to the ecology of the genus in Russia.

*Testechiniscus* distribution worldwide follows a similar pattern: reports confirmed with molecular barcoding show that *T. spitsbergensis* gravitates towards the northern Atlantic (Greenland, Scotland, Norway and Spitsbergen as terra typica), as well as central Europe (Poland). Recent morphological records show its presence in Ireland (DeMilio et al. 2016). The subspecies *T. s. tropicalis* is reported from a mountainous region in Ethiopia, which can be considered an exception to *Testechiniscus* geographical distribution pattern (Gąsiorek et al. 2018b). Finally, *T. laterculus* has been recorded from Canada (Massa et al. 2024), with its type locality being Riverton in northern California, United States (Schuster & Grigarick 1965).

## SUPPLEMENTARY DATA

Electronic supplementary material 1. Primers and PCR programs used for amplification of four DNA fragments sequenced in the study.

Electronic supplementary material 2. Complete list of sequences used in the molecular phylogenetic analysis.

Electronic supplementary material 3. Video demonstrating claw movements of the new species. Format: MP4

## CONFLICT OF INTEREST

The authors declare that they have no conflicts of interest in relation to this study.

## ACKNOWLEDGEMENTS

We are grateful to Arseniy Kudikov (Russian Academy of Sciences) for collecting samples from Novaya Zemlya archipelago, Boris Kochnev (Novosibirsk State University) for collecting samples from Krasnoyarskiy kray, and to Vadim Panov for collecting samples from Ireland. This study was carried out with the use of the equipment of the Core Facilities Centre “Centre for Molecular and Cell Technologies” of St Petersburg State University (https://researchpark.spbu.ru/index.php/en/biomed-eng) and “Taxon” Research Resource Center of the Zoological Institute of the Russian Academy of Sciences (http://www.ckp-rf.ru/ckp/3038/).

## FUNDING

The study was supported by the Russian Science Foundation, grant No. 25-74-20033 “Evolutionary transformations of nanostructural elements of the Ecdysozoa cuticle using the example of the integumentary structures of tardigrades (Tardigrada)”. D.V.T. received support for the field trip to Sakhalin from non-profit charitable foundation “Support of bioresearch “BIOM”, grant № 4/2024-гр.

## DATA AVAILABILITY STATEMENT

The data underlying this article will be shared on reasonable request to the corresponding author.

## Notes

### Competing Interest Statement

The authors have declared no competing interest.

