## Supplementary Material 01 for "Integrative description of a new species of *Testechiniscus* (Tardigrada, Echiniscoidea) from Novaya Zemlya, Russia, with discussion of the genus diagnosis and distribution"

**Primers and PCR programs used for amplification of the four DNA fragments sequenced in the study.**

**Electronic supplementary material 1 to the article:**

| DNA fragment | Primer name | Primer direction | Primer sequence (5’–3’) | Primer source | PCR programme |
| --- | --- | --- | --- | --- | --- |
| COI | LCO1490-JJ | forward | CHACWAAYCATAAAGATATYGG | Astrin and Stüben, 2008 | Michalczyk et al., 2012 |
|  | HCO2198-JJ | reverse | AWACTTCVGGRTGVCCAAARAATCA |  |  |
| 18S rRNA | 18S_Tar_Ff1 | forward | AGGCGAAACCGCGAATGGCTC | Stec et al., 2017 | Zeller (2010), in Stec et al., 2015 |
|  | 18S_Tar_Rr1 | reverse | GCCGCAGGCTCCACTCCTGG |  |  |
| 28S rRNA | 28S_Eutar_F | forward | ACCCGCTGAACTTAAGCATAT | Gąsiorek et al., 2018 | Mironov et al.*,* 2012 |
|  | 28S_R0990 | reverse | CCTTGGTCCGTGTTTCAAGAC | Mironov et al., 2012 |  |
| ITS-1 | ITS1-Echi-F | forward | CCGTCGCTACTACCGATTGG | Gąsiorek et al., 2018 | Stec et al., 2018 |
|  | ITS1-Echi-R | reverse | GTTCAGAAAACCCTGCAATTCACG |  |  |
