## Supplementary Material 02 for "Integrative description of a new species of *Testechiniscus* (Tardigrada, Echiniscoidea) from Novaya Zemlya, Russia, with discussion of the genus diagnosis and distribution"

**Complete list of sequences used in this study. Sequences produced in this study are marked in bold.**

**Electronic supplementary material 2 to the article:**

| Species | Locality | 18S rRNA | 28S rRNA | ITS-1 | COI |
| --- | --- | --- | --- | --- | --- |
| *Testechiniscus spitsbergensis spitsbergensis*  Scourfield, 1897 | Spitsbergen, Norway | MH279664  MH279665 | MH286186  MH286188 | MH286189 | - |
|  | Krasnoyarskiy kray, Russia | - | - | - | PX506518 |
|  | Sakhalin, Russia | - | - | - | PX506519  PX506520 |
|  | Disko Island, Greenland | EU266967 | - | - | HM193419  JX676199 |
| *Testechiniscus spitsbergensis tropicalis* Gąsiorek et al., 2018 | Bwahit, Ethiopia | MH279666 | MH286187 | MH286191 | - |
| *Testechiniscus laterculus* (Schuster, Grigarick and Toftner, 1980) | British Columbia, Canada | OQ029311 | - | - | OQ029483 |
| *Testechiniscus impenetrabilis* **sp. nov.** | Novaya Zemlya, Russia | PX584498  PX584499  PX584500  PX584501 | PX584502  PX584503  PX584504  PX584505  PX584506 | PX584446  PX584447  PX584448  PX584449  PX584450 | PX506521  PX506522  PX506523  PX506524  PX506525 |
